# Leveraging Soil Mapping and Machine Learning to Improve Spatial Adjustments in Plant Breeding Trials

**DOI:** 10.1101/2024.01.03.574114

**Authors:** Matthew E. Carroll, Luis G. Riera, Bradley A. Miller, Philip M. Dixon, Baskar Ganapathysubramanian, Soumik Sarkar, Asheesh K. Singh

## Abstract

Spatial adjustments are used to improve the estimate of plot seed yield across crops and geographies. Moving mean and P-Spline are examples of spatial adjustment methods used in plant breeding trials to deal with field heterogeneity. Within trial spatial variability primarily comes from soil feature gradients, such as nutrients, but study of the importance of various soil factors including nutrients is lacking. We analyzed plant breeding progeny row and preliminary yield trial data of a public soybean breeding program across three years consisting of 43,545 plots. We compared several spatial adjustment methods: unadjusted (as a control), moving means adjustment, P-spline adjustment, and a machine learning based method called XGBoost. XGBoost modeled soil features at (a) local field scale for each generation and per year, and (b) all inclusive field scale spanning all generations and years. We report the usefulness of spatial adjustments at both progeny row and preliminary yield trial stages of field testing, and additionally provide ways to utilize interpretability insights of soil features in spatial adjustments. These results empower breeders to further refine selection criteria to make more accurate selections, and furthermore include soil variables to select for macro– and micro-nutrients stress tolerance.

## 1 Introduction

Plant breeders make selection decisions within their programs to advance lines with the highest genetic value for the target population of environments [1]. Breeders must address non-uniform field conditions (i.e., field heterogeneity) in the selection decision making process, as an incorrect decision causes economic strain on the program and limits success [2]. Spatial adjustment methods have been proposed to alleviate the challenges of non-uniform field conditions. These methods allow breeders to make more appropriate comparisons of entries to each other, as well as to checks in yield plot testing. Spatial adjustments set up an effective and efficient selection process in the plot testing stages. Spatial adjustments are particularly applicable in unreplicated trials such as early stage yield testing of hybrids in cross-pollinating crop species, and for purelines in progeny row (PR) and preliminary yield trials (PYT) stages in self-pollinating crops.

Reducing the size of the error associated with each genotype (i.e., pureline) gives breeders the ability to discriminate between yield levels of competing genotypes for selection more accurately; and therefore several methods have been proposed. Augmented designs (AD) with replicated checks [3], give some form of local control within a field, giving the breeder a reasonable estimate of the trends that unreplicated genotypes may be experiencing. AD offers breeders the possibility of estimating the standard errors based on the replicated check performance between similar counterparts. Check plot methods can be used by placing checks throughout the field so that a breeder can adjust for field trends based on known checks and their relative performance [4]. An advantage of the check plot method is that there is no limit to the number of lines used in the trial; on the other hand, it requires a large number of experimental plots to be allocated to the check plots [4]. The grid method was introduced for a mass selection experiment in maize [5], where small grids of 40 plants across the field were set up and the top-yielding 10% of plants in each grid were selected to minimize the spatial trends within the field. Another popular method is the moving means (MM) method [6], where the use of a MM coefficient to adjust yields resulted in a significant decrease in the error when estimating genotypic effects. Most spatial adjustment methods correct for field trends and do not directly quantify the soil features for each plot.

A common theme in all spatial adjustment experiments for yield trials is the axiom that these methods reduce the effects of soil heterogeneity, thereby facilitating an unbiased comparison of plot values. While there is a general acceptance of this notion in breeding trials, limited or no information is available that directly quantifies the effects of different soil characteristics and use these values to adjust plot yields. Soil is an integral feature of all breeding testing, with scant consideration of accounting for within field variability. Soil features include calcium, potassium, magnesium, phosphorous, cation exchange capacity, organic matter, pH, sand, silt and clay. Instead of directly quantifying soil variables and their effect on yield, the spatial adjustment methods either model the yield trends or reduce the area of the selection into smaller homogeneous blocks. These methods reduce the standard error of the difference between estimated genotypic values that are being evaluated, and increases the precision in which yield differences can be compared within breeding trials [7].

The utilization of soil features can provide an association between genotype, soil, and yield opening up avenues for improvement and utilization of spatial adjustments. Despite the use of soil characterization in production fields [8–10], they have not been utilized in breeding trials due to the resolution needed for soil maps, as well as the complexity of parsing out environmental and genotypic variance within a model. Furthermore, in breeding trials, the scale of experiments (# of entries, locations, years etc.) necessitates machine learning (ML) approaches to generate relationships and learn trends along with improved interpretability.

Machine learning is a powerful tool in plant science research, with wide ranging trait investigations such as biotic identification and quantification[11–21], organ detection [22–26], microscopic objects [27, 28], abiotic stresses [29–32], and crop yield [33–35]. These ML methods provide an opportunity to integrate multiple variables for spatial adjustment while parsing out the trends and role of soil features on plot yield in tests. Among the extensive diversity of methods in ML, “Extreme Gradient Boosting” (XGBoost) [36] is well suited for the spatial adjustment problem since it is a hierarchical ensemble modeling tree structure that derives its outputs utilizing classification rules. XGBoost is scalable and gives high model accuracy, allows for utilization of multiple features in prediction, and has been shown to be very useful in regression and classification problems [36, 37]. XGBoost, like other decision tree methods, is an algorithm that can effectively use multiple variables included in a dataset for prediction. Decision tree structures do not assume a linear relationship between variables, or the predicted values, which can help to predict nonlinear multivariate problems. This structure allows for variables to have multiple weights depending on the values of other variables. This is particularity applicable to soil plant growth interactions, as it is known that plant growth and development is dependent on multiple interacting soil features, weather, and genetics.

The motivation for the study was to create a paradigm shift from current spatial adjustments of yield plot data, primarily made on the plot yield values of neighbors, to a method that includes plot yield and soil features. To achieve our goals, digital soil maps and ML were leveraged to model the effect of soil features on plot yield in a soybean breeding program. XGBoost made it possible to adjust for the growing conditions experienced by each genotype. Furthermore, we present the use of Shapley values to aid in the model interpretability and explore its usefulness to breeders by providing previously underutilized knowledge about the soil conditions from trials, and its integration in breeding decisions. Finally, we propose the usefulness of this method to consciously select for nutrient deficiency tolerance in field trials that experience abiotic stresses.

## 2 Materials and Methods

### 2.1 Field experiments and data collection

Soybean [*Glycine Max (L.) Merr.*] breeding trial plot data were collected in three growing seasons (2019, 2020, 2021). Plot yield data were from progeny rows (coded as A-test; 2019, 2020, 2021) and next stage testing, i.e., preliminary yield trials (coded as B-test; 2019, 2020, 2021). Data was collected from the central Iowa research locations, centered around the Ag Engineering and Agronomy research farm (Boone, IA, USA). A brief description of each field can be found in table 1, no border rows were used between plots. The centers of plots in the same row were 1.52 meters apart. Plot lengths are recorded as the harvested length of each plot, all fields had a .91 meter alleyway between plots in each column giving a plot to plot center distance of 3.04 and 6.09 meters for plots in the same column for the A and B tests, respectively. To minimize interplot competition, each field consisted of multiple experiments that grouped lines with similar crop maturity, phenology and genetic background. Plots were harvested with an ALMACO (Nevada, IA USA) small plot combine, and yields were adjusted to 13% moisture, and converted to kilograms/hectare (kg/ha) for each harvested plot.

**Table 1:** Listing of A-test (i.e., progeny row tests), and B-tests (i.e., preliminary yield trials) that were used in this study. Tests in each year, location, geographical location, and plots details are listed.

| Year | Test | Location | Latitude | Longitude | Number of Plots | Avg. Replicated Check Count | Number of Checks | Experiment Count | Plot Rows | Row Spacing (m) | Plot Length (m) | Plot Area ($m^2$) | Planting Density (100k seeds/hectare) |
| --- | --- | --- | --- | --- | --- | --- | --- | --- | --- | --- | --- | --- | --- |
| 2019 | A | Marsden | 42.010N | 93.780W | 10870 | 77.9 | 10 | 53 | 2 | 0.76 | 2.13 | 3.25 | 346 |
| 2019 | B | Marsden | 42.010N | 93.778W | 3062 | 30.8 | 10 | 21 | 2 | 0.76 | 5.18 | 7.90 | 346 |
| 2020 | A | AEA | 42.015N | 93.768W | 1458 | 25.714 | 7 | 8 | 2 | 0.76 | 2.13 | 3.25 | 346 |
| 2020 | A | Burkey | 42.012N | 93.788W | 3806 | 24.545 | 11 | 19 | 2 | 0.76 | 2.13 | 3.25 | 346 |
| 2020 | A | Burkey.CH | 42.015N | 93.792W | 2920 | 33.4 | 10 | 13 | 2 | 0.76 | 2.13 | 3.25 | 346 |
| 2020 | B | CAD | 42.047N | 93.739W | 3272 | 33.75 | 12 | 24 | 2 | 0.76 | 5.18 | 7.90 | 346 |
| 2021 | A | Marsden.East | 42.011N | 93.778W | 11644 | 92.5 | 12 | 54 | 2 | 0.76 | 2.13 | 3.25 | 346 |
| 2021 | A | Marsden.West | 42.010N | 93.781W | 3479 | 60.4 | 5 | 16 | 2 | 0.76 | 2.13 | 3.25 | 346 |
| 2021 | B | Burkey | 42.015N | 93.788W | 3034 | 29.833 | 12 | 30 | 2 | 0.76 | 5.18 | 7.90 | 346 |

### 2.2 Soil data collection

Soil data was collected on a 25 meter grid pattern for each field at a coring depth of 15cm, and then sent to Midwest labs (Omaha, NE USA) for analysis. Soil maps were created from the 25 meter grid samples to a resolution of 3×3 meter grids for soil nutrients and particle size fractions [Calcium (Ca), Cation Exchange Capacity (CEC), Phosphorous (P1), soil pH (PH), Magnesium (MG), Potassium (K), Organic Matter (OM), Clay, Silt, and Sand], using the Cubist machine learning algorithm [38]. Soil values were extracted for each plot using Esri’s ArcGIS Python library (Redland CA, USA), arcpy, and Zonal Statistics toolbox. The extracted soil values for each plot were then used to calculate mean soil values on a per plot basis using a 3×3 grid, which is represented by Eq. 1. Where 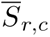 is the mean soil feature value of the *r^th^* row and *c^th^* column in each field. All further analysis using soil features used the 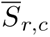 values as input.

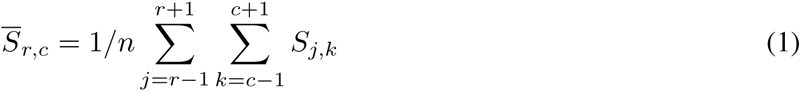

### 2.3 Base Line Spatial Adjustments

#### 2.3.1 Moving Means

Plot seed yields were adjusted using Moving Means with a grid size of 5×5 plots, and were calculated using the R package “mvnGrad” [39]. This method excluded the plot of interest when calculating the mean for each plot. The Moving Means for each experiment within a test were calculated separately. The Moving Means method makes an assumption that neighboring plots do not exhibit interplot competition, and that the nonrandom error observed in a field is due to field environmental trends [40]. Breeders can reduce the interplot competition by subdividing fields into smaller experiments that are grouped by similar maturities and genetic backgrounds.The process for calculating the Moving Means can be found in the mvnGrad documentation [39]. Results from the Moving Mean model from here on will be referred to as ADJ 5 MM.

#### 2.3.2 P-Splines

P-splines are an alternative method to Moving Means, commonly used in spatial adjustments for field trials. The R package “statgenSTA” [41] was used to correct for field variations using P-splines as described in Rodriguez et al. [42]. For this study, genotypes were treated as a random factor. The design was specified as a row-column design, which treats the row and column as random effects, and used the “SpATS” engine. Results from the P-Spline model from here on will be referred to as Spline.

### 2.4 Soil-Based Adjustments

#### 2.4.1 Statistical Learning

XGBoost training process requires tuning parameter coefficients, usually denoted by *θ*. Among all the parameters, the number of estimators, maximum depth, learning rate, *γ*, subsample size, and minimum child weights are commonly tuned using a greedy search algorithm. Most XGBoost models are configured using a relatively shallow depth or a small number of trees because of their sequential characteristic, where each new tree corrects the errors made by previous trees, quickly reaching a point of diminishing returns. This training structure works well for soil-based spatial adjustments, because it is able to train models quickly, and model complex data with multiple interactions. Our model used 240 estimators and a depth of 9 for training. Model parameter tuning was done using *Sklearn grid search API* [43] with 10-fold cross validation. Two methods were compared for spatial adjustments using XGBoost. Model 1, from here on referred to as XGB Global, combined data from all nine fields and attempted to make a generalized model. Model 2, from here on referred to as XGB Local, was trained on each field individually to create a more specific model for each field location. Both models used the same tuning parameters as listed in table 2.

**Table 2:**
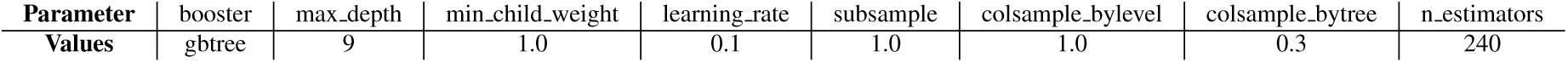
Parameters used for the XGBoost model.

#### 2.4.2 Model overview

Plant breeders routinely express yield as *P* = *G* + *E*, where *P* is the observed phenotypic value of a line, e.g., seed yield. *G* is the genotypic value of the line, which can be estimated when lines are replicated and grown in multiple locations. Lastly, *E* is the environmental effect, which breeders, try to minimize, as it can confound the estimated genetic value of a line. Historically, *E* has been a macro-scale environment effect such as a single field, and micro-scale plot level environmental effects have been reduced using various spatial adjustment methods. Furthermore, genotypic value can be better estimated by using phenotypic values of related lines. If the mean of the population is included, the new equation is represented as *P* = *G* + *E* + *µ*, where *µ* is the phenoytpic mean of the population. In this context, we define population to consist of purelines from the same breeding family, that is they have the same the pedigree. Our proposed method uses soil features to estimate the value of *E* for each plot, and to increase the correlation between the adjusted phenotypic values and the estimated genotypic values.

The first step was to collect the yield data for each plot in a given field which was in a row and column configuration within each field. Each experiment has multiple lines that are from the same or related populations, and the mean of each population within each experiment was calculated and used as the estimate for the population mean within the experiment. Checks were averaged across the entire field and this average was used as the estimate for the check population mean. Eq. 2 represents the basis for the soil-based adjustment methods. *P_r,c,j,k_* is the the seed yield of the plot in the *r^th^* row of the *c^th^* column from the *j^th^* population in the *k^th^* experiment. *G_r,c,j,k_* is the the genetic value of the plot in the *r^th^* row of the *c^th^* column from the *j^th^*population in the *k^th^* experiment. *S_r,c_* is the soil feature values of the plot in the *r^th^* row of the *c^th^* column. *µ_j,k_* is the mean of all plots in the *j^th^* population of the *k^th^* experiment.

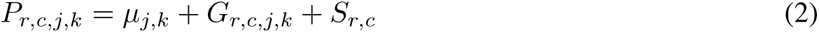

The next step was to subtract the population mean within each experiment from the observed yield of each plot, giving the phenotypic deviation, *δ_r,c_*, from the population mean of each plot, this is shown in Eq. 3. It is assumed that part of *δ_r,c_* can be attributed to the soil features and part can be attributed to the genetic value of the line.

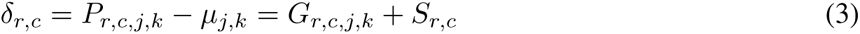

The third step was then to train a model, using XGBoost regression, to predict the deviation, *δ_r,c_*, from the population mean based on the soil features, *S_r,c_*, of each plot. This trained model was then used to estimate the effect of the soil on each plot’s observed yield, represented by 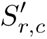 in Eq. 4

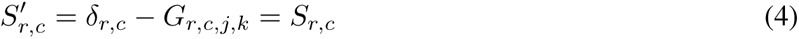

The last step estimates the genetic value of each plot by taking the observed phenotypic value, and subtracting the estimated effect of the soil, 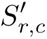, from the observed phenotypic value, *P_r,c,j,k_*, which is represented as 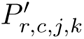 in Eq. 5. 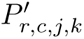 was used for further analysis and comparison to other spatial adjustment techniques.

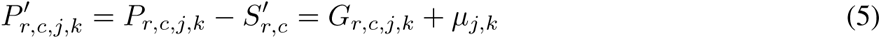

### 2.5 Model Interpretability

‘SHAP” (SHapley Additive exPlanations) [44] was used to assist interpreting the results from XGBoost models. SHAP uses traditional Shapley values from game theory, and it has shown to be very successful in explaining the output of any machine learning model [45, 46]. Shapley values estimate the feature’s importance in terms of their contribution to the outcome.

### 2.6 Model Performance for Selection

Three evaluation metrics were used to assess the performance of the different methods. The first metric was the relative efficiency using the average standard error of the difference (*SED*) between the check lines in each field. The efficiency of different spatial adjustment methods was calculated using Eq. 6 where *SED_Y ield_* is the *SED* using no spatial adjustments, and *SED_Adj_*is the *SED* using one of the tested spatial adjustment methods.

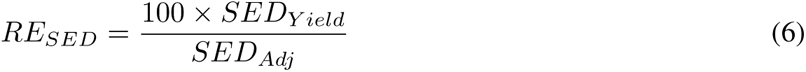

In A-test, entries are not replicated due to seed limitation and in B-test stage breeders prefer to test in more than one location in unreplicated manner; therefore, checks were used to estimate the standard error of difference. Previous works evaluating the efficiency of spatial and experimental designs in plant breeding trials have used this as a method for evaluation of different selection methods [7, 47, 48]. As the standard error of the difference decreases, more of the observed phenotypic yield variance can be attributed to the genotypic value. The number of genetically distinct checks and the average number of times each were replicated in the field can be found in table 1.

The second metric used to evaluate the models was a similarity coefficient called the Czekanowski coefficient (*CZ*), also known as the Sorensen Dice coefficient. This coefficient gives the percent of lines selected by both selection methods at a given selection intensity. The Czekanowski coefficient was calculated according to [7] where *a* is the number of lines selected by both selection methods, *b* is the number of lines selected only by model 1, and *c* is the number of lines selected only by model 2. *CZ* is a ratio given by Eq. 7. If *CZ* has a value of 1 it would mean that both methods selected the same lines, if *CZ* has a value of 0 it would mean that there was no overlap in the lines that were selected. For our purposes model 1 used no spatial adjustment, and model 2 was one of the listed spatial adjustment methods.

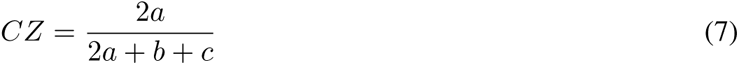

The last metric is the Moran’s I statistic, a measure of the spatial autocorrelation present within a user defined grid [49]. Our implementation uses a grid of 5 by 5 plots to define the plot neighbors given the row column field design of each test. Neighbor distances used relative coordinates to determine plot neighbors, plot widths and lengths were considered to have a unit of 1. Plots that were within a distance of 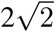 of a plot were considered neighbors. All other plots were not considered to be neighbors. Moran’s I interpretation is similar to a regular correlation coefficient with values ranging from approximately –1 to 1. A value of 1 is a perfect spatial autocorrelation, and a value of –1 is a negative spatial autocorrelation. The null hypothesis expected values for the Moran’s I statistic are close to zero, but are slightly negative. P-values are also reported with each statistic to assist in the interpretation of the significance of the correlation. Moran’s I statistics were calculated in R version 4.0.3 with the package “spdep” [50] and the Moran.test function.

## 3 Results

### 3.1 Genetic Variation

Across three years, 238 experiments were examined across nine individual fields. Experiment sizes ranged from 50 to 370 plots with a mean of 183.0 plots per experiment. The A and B-test plot number means were 209.7 and 124.9 plots respectively. In these 238 experiments, 282 unique breeding populations were assessed. Breeding population sizes ranged from 1 to 263 genotypes with a mean size of 70.6. The average population size in the A and B test, was 133.9 and 42.3, respectively. Population mean seed yields ranged from 74.0 to 6032.3 kilograms per hectare (kg/ha) with a mean yield of 4250.2 (kg/ha). The average population seed yields for the A and B tests were 4391.4 and 3624.8 (kg/ha) respectively. While B-tests have one additional generation of selection in the previous year, the lower seed yield is likely due to shorter plot size in A-test (2.3 m) compared to B-test (5.18 m) giving a larger plot edge effect [51]. Table 3 gives a further summary of the data across all nine fields.

**Table 3:** Listing of the fields evaluated for this study. The number of experiments, average experiment plot number, mean population size, and mean population seed yields are listed. Values in parentheses are the standard deviations.

| Field | Experiment Count | Mean Number of Plots | Number of Populations | Mean Population Number | Mean Population Yield (kg/ha) |
| --- | --- | --- | --- | --- | --- |
| 2019_A_Marsden | 53 | 205.1 (54.5) | 60 | 162.9 (47.1) | 3819.8 (383.3) |
| 2019_B_Marsden | 21 | 145.8 (53.4) | 53 | 45.7 (20.3) | 3160.8 (295.9) |
| 2020_A_AEA | 8 | 182.2 (56.9) | 55 | 22.0 (17.1) | 3914.0 (847.4) |
| 2020_A_Burkey | 19 | 200.3 (71.8) | 23 | 149.2 (58.3) | 4438.5 (766.7) |
| 2020_A_Burkey_CH | 13 | 224.6 (54.9) | 43 | 56.5 (34.3) | 3732.4 (578.4) |
| 2020_B_CAD | 24 | 136.3 (53.3) | 61 | 43.1 (22) | 3107.0 (423.7) |
| 2021_A_Marsden_East | 54 | 215.6 (67.5) | 90 | 117 (27.7) | 5124.5 (376.6) |
| 2021_A_Marsden_West | 16 | 217.4 (59.2) | 90 | 35.1 (15.5) | 5285.9 (417.0) |
| 2021_B_Burkey | 30 | 101.1 (31.6) | 64 | 38.8 (28.8) | 4505.8 (430.4) |

### 3.2 Soil Variability

Soil core samples were taken for each of the nine fields with ten soil features measured (see M and M). Organic matter ranged from 1.1% to 7.4% with a mean of 3.9%. PH ranged from 3.9 to 8.5 with a mean of 6.1. Clay had a range from 0.1% to 33.9% with an mean of 27.4%. Sand had a range from 12.2% to 42.7% with a mean of 30.4%. Silt had a range from 18.9% to 50.5% with a mean of 41.6%. Calcium parts per million (ppm) ranged from 507.1 to 6360.5 with a mean of 2774.9. CEC (meq/100 g soil) ranged from 9.8 to 36.5 with a mean of 21.1. Potassium (ppm) had a range from 85.4 to 286.1 with a mean of 169.1. Magnesium (ppm) ranged from 107.3 to 768.9 with a mean of 399.7. Phosphorous (ppm) had a range from 5.5 to 73.5 with a mean of 21.7. Table 4 shows the range of each soil parameter for each individual field to better represent the variation that was present across locations. It should be noted that due to a small sized 2020 A AEA field test, there was a lack of variability in the samples taken in the clay content, and this value was constant for that test.

**Table 4:** Soil descriptive statistics for each field. The minimum, maximum, and mean values are listed.

| Field |  | MM_CA | MM_CEC | MM_K | MM_MG | MM_OM | MM_P1 | MM_PH | MM_CLAY | MM_SAND | MM_SILT |
| --- | --- | --- | --- | --- | --- | --- | --- | --- | --- | --- | --- |
| 2019_A.Marsden | min | 507.11 | 9.78 | 109.33 | 107.33 | 1.07 | 8.78 | 3.88 | 24.77 | 20.33 | 37.42 |
|  | mean | 2262.71 | 18.95 | 157.07 | 323.62 | 3.34 | 21.84 | 5.9 | 28.34 | 29.12 | 46.38 |
|  | max | 4918.78 | 33.11 | 193.33 | 768.89 | 5.58 | 37.33 | 8.5 | 33.82 | 39.19 | 42.32 |
| 2019_B.Marsden | min | 1147.11 | 11.89 | 110.5 | 158.33 | 1.71 | 9.11 | 4.26 | 22.55 | 21.13 | 38.9 |
|  | mean | 2543.17 | 20.79 | 160.56 | 379.01 | 3.97 | 18.66 | 6.22 | 27.29 | 29 | 43.49 |
|  | max | 4986 | 36.5 | 218.33 | 754.56 | 6.5 | 36.33 | 8.3 | 30.75 | 37.87 | 49.45 |
| 2020_A.Burkey | min | 1748.56 | 12 | 133 | 253.67 | 2.15 | 11.22 | 4.78 | 14.1 | 15.64 | 20.53 |
|  | mean | 2799.29 | 20.75 | 179.57 | 390.96 | 3.62 | 25.25 | 6.05 | 26.5 | 32.76 | 38.24 |
|  | max | 3818.5 | 27.83 | 212.5 | 594.11 | 5.13 | 73.5 | 7.23 | 33.9 | 42.73 | 40.1 |
| 2020_A.Burkey.CH | min | 2212.44 | 14 | 85.44 | 218.5 | 2.61 | 6.44 | 6.1 | 21.68 | 30.79 | 18.9 |
|  | mean | 2810 | 18.58 | 152.16 | 318.33 | 3.29 | 14.9 | 6.44 | 25.7 | 35.96 | 37.4 |
|  | max | 4061.83 | 27 | 286.11 | 578 | 4.6 | 47.17 | 6.9 | 30.08 | 41.75 | 39.5 |
| 2020_A.AEA | min | 2012.83 | 15 | 127.78 | 239.83 | 3 | 6.33 | 5.8 | 28.9 | 29.07 | 37.5 |
|  | mean | 3315.97 | 21.89 | 1551.9 | 412.69 | 3.85 | 14.08 | 6.54 | 28.9 | 30.72 | 40.33 |
|  | max | 4040.67 | 26.56 | 170.33 | 526 | 4.3 | 33 | 7.7 | 28.9 | 32.76 | 41 |
| 2020_B.CAD | min | 1632.5 | 12.33 | 134.67 | 206 | 2.51 | 5.5 | 5.67 | 0.08 | 12.16 | 32.97 |
|  | mean | 3829.95 | 23.89 | 183.19 | 358.08 | 4.56 | 13.93 | 6.8 | 25.25 | 31.64 | 40.18 |
|  | max | 6360.5 | 35.83 | 236 | 496.44 | 6.79 | 27.56 | 8.2 | 33.6 | 39.27 | 43.54 |
| 2021_B.Burkey | min | 1748.85 | 14.85 | 153.28 | 202.56 | 2.86 | 9.34 | 5.49 | 22.13 | 27.98 | 35.17 |
|  | mean | 2606.55 | 20.58 | 177.45 | 361.54 | 3.8 | 21.46 | 5.77 | 25.94 | 35.11 | 38.95 |
|  | max | 3732.62 | 26.77 | 217.52 | 555.57 | 4.9 | 32.02 | 7.12 | 29.84 | 42.24 | 42.93 |
| 2021_A.Marsden.West | min | 1997.02 | 14.99 | 135.46 | 633.28 | 2.62 | 14.33 | 5.51 | 25.08 | 23.22 | 37.37 |
|  | mean | 2644.48 | 21.28 | 170.37 | 456.05 | 3.88 | 28.26 | 5.96 | 28.4 | 29.98 | 42.38 |
|  | max | 4281.7 | 28.7 | 217.26 | 633.28 | 5.5 | 39.97 | 6.82 | 31.79 | 39.04 | 46.29 |
| 2021_A.Marsden.East | min | 1717.1 | 15.76 | 119.81 | 312.63 | 2.64 | 8.87 | 4.92 | 22.14 | 19.2 | 37.47 |
|  | mean | 3015.68 | 22.96 | 179.16 | 502.15 | 4.44 | 24.16 | 6.09 | 27.69 | 28.41 | 43.71 |
|  | max | 5896.5 | 34.19 | 228.69 | 733.77 | 7.37 | 43.14 | 7.89 | 32.62 | 40.47 | 50.53 |

### 3.3 XGBoost and Model Interpretability

XGB Local model’s predictions, ground truth and predicted yield histograms, and correlation are shown in Fig. 1. These predictions are on the centered variables, as described in Eq. 3 as *δ_r,c_*. The Concordance Correlation Coefficient between the ground truth and predicted values ranged from 0.71 to 0.86. Centered values, ranged from –81.5 to 48.8. When looking at the two stages of the breeding program, i.e., progeny row A-test and preliminary yield trial B-test, *δ_r,c_* values in A-tests ranged from from –81.5 to 48.8 and in B-tests ranged from –47.7 to 32.5. This demonstrates that the more advanced stage of testing has less phenotypic variability. Predicted values, which were obtained from XGB Local model and reflect estimated effects of soil features on seed yield, ranged from –62.7 to 36.3. These values represent *S^′^* as defined in Eq. 4. The A-test predicted values ranged from –62.7 to 36.3 and the B-test predicted values ranges from –40.2 to 21.6.

**Figure 1:**
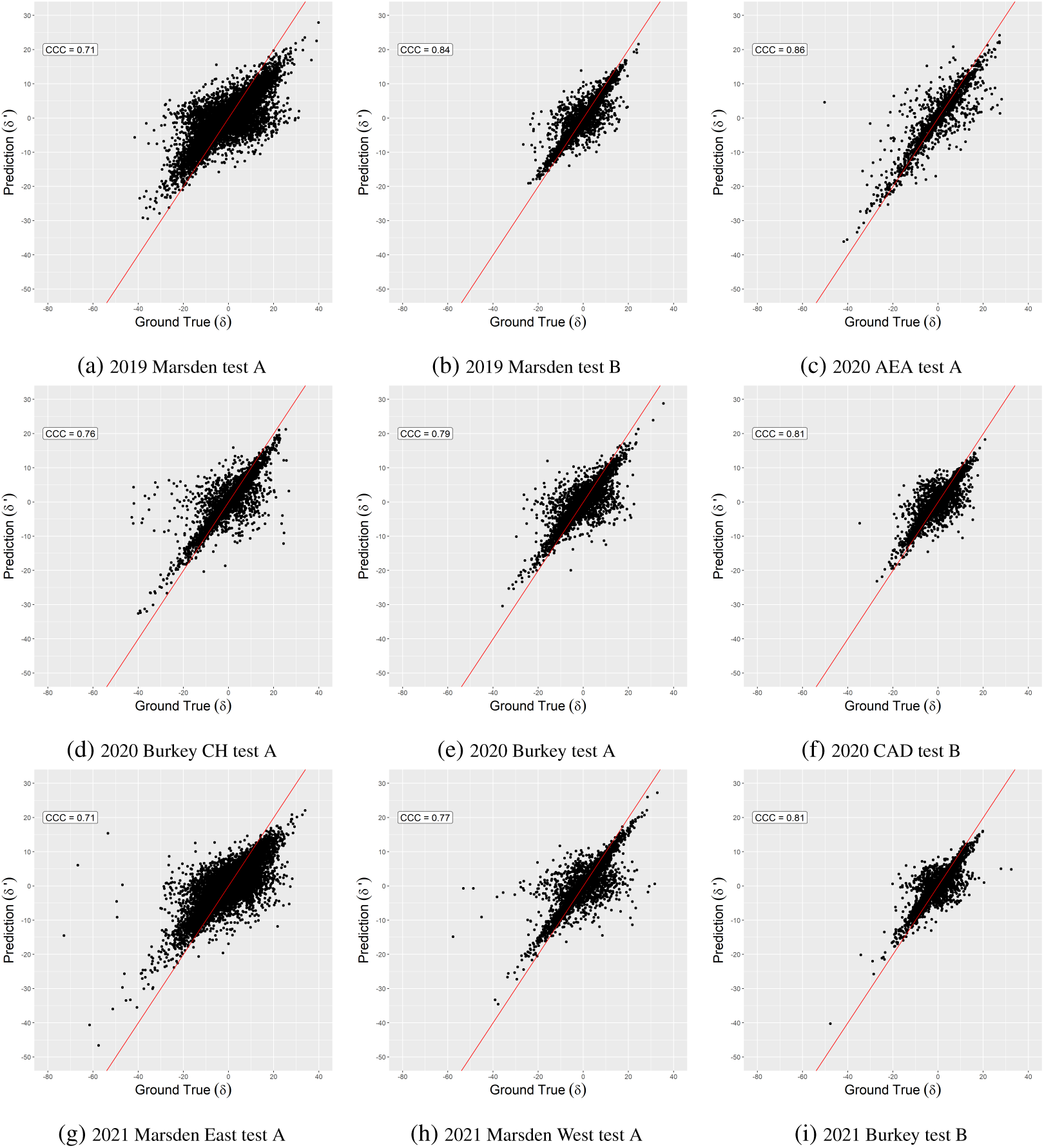
This figure shows the XGB Local Ground true (*δ*) vs prediction (*δ^′^*) scatter plots with their respective Concordance Correlation Coefficient. These charts show how well the model learned the ground true data distribution. The red line shows a 1:1 relationship between predicted and observed values

Fig. 2 shows results as an example for the 2021 A Marsden East test. In feature selection, correlation provides a measure of independence between input variables; the higher the correlation, the greater the linear dependency. The heatmap plot in Fig. 2a shows the correlation between the features, and the dendrogram on top provides a clustering mechanism of these features according to their correlation values. The inverse “u” shape-like links group elements into clusters; the lower the “u” shape is in the figure, the higher the correlation of those elements is in relation to the others. The MM OM and MM CEC correlation is 0.9, the highest between elements, and are clustered in the dendrogram first. Then the MM OM MM CEC cluster is merged with MM CA to form another cluster. For this article, highly correlated features were kept to explore their outcomes using Shapley values.

**Figure 2:**
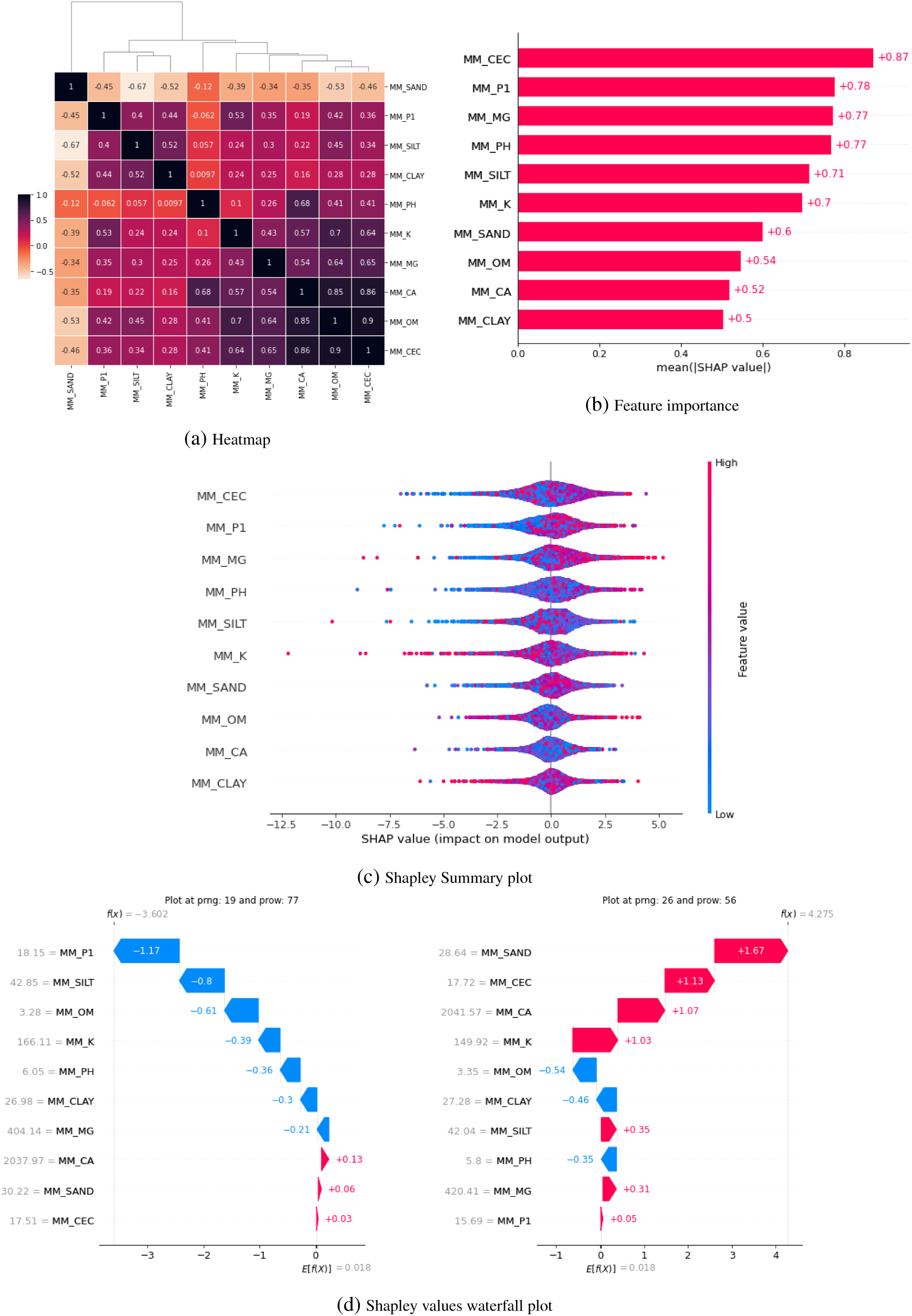
XGBoost model explainability plots using data from 2021 Marsden East Test A. The heatmap (a) shows the correlation between the features used to train the XGBoost model and their respective clustering hierarchy by the dendrogram. The feature importance, figure (b), illustrates a measure of the prediction power of each feature for the model with this dataset. Each dot in the Shapley value Summary plot (c) represents a single plot. Its colors are normalized per feature using its maximum and minimum respectively. The summary plot helps illustrate the relationship between the individual feature value range and their impact over the predicted yield. The Shapley values waterfall plots (d) provide an insight into how features values contribute to the final expected value in two plots (Identities “IA2102”) at different locations in the field where the feature values are substantially different.

Fig. 2b provides a summary of the features of importance for the model. The higher the value on the bars in this plot, the greater the information gained by the model when performing the tree branch split during training. In the 2021 Marsden A test East, MM CEC provided the most significant information gain, and the other top four driving prediction features were MM P1, MM MG, and MM PH.

To aid in interpreting results we investigated the use of Shapley values. Fig. 2c describes how each plot reacted to the input features. Each dot is a plot, and its color represents the plot feature value relative to the feature’s range scale, where red is high and blue is low. The accumulation of the dots across the x-axis characterizes the global behavior of the features, similar to a histogram. For the 2021 Marsden A test East, we see that high values of MM CEC, MM P1, and MM MG tend to have positive Shapley values. However, it is not conclusive for the other soil features, where we see high feature values on both tails. For our model, we can not draw any causal inference conclusions because we suspect the existence of many confounding factors.

The waterfall plots in Fig. 2d describe the reaction of the the check genotype (IA2102) compared to the soil composition in distinct locations of the field used on the 2021 Marsden A test East. A check variety was used so that the comparison is only looking at the soil features and not including additional genetic differences between two lines. The model’s mean output prediction for the entire field is 0.018. If we start from the mean predicted value of 0.018 and add the respective Shapley values, we obtain the model prediction of 4.275 for the right image and –3.602 for the left. This represents the relative quality of the growing conditions for each individual plot, and the expected contribution of each soil feature, as determined by the model.

### 3.4 Spatial Adjustments

Mean relative efficiencies were calculated for each model, with values of 112.8%, 132.8%, 161.6%, and 178.6% for ADJ 5 MM, XGB Global, Spline, and XGB Local. No spatial adjustments was the base model to which all models were compared. The models are ranked from lowest to highest relative efficiency; where the Moving Mean method has the lowest relative efficiency and XGB Local has the highest average relative efficiency across all nine fields that were tested. All of the models presented have a higher relative efficiency than the unadjusted yields. This indicates that XGBoost using plot level soil fertility variables can be used to increase the precision of the genotypic estimates of the checks. These results infer that the estimated genotypic values for unreplicated lines will be more precise as well. Increasing the precision of the estimates of the genetic values of a line will lead to higher selection accuracy and increases in genetic gain.

Similarity coefficients were calculated at three selection intensities (0.1, 0.3, and 0.4), and examined the differences in the selected lines based on the method used for spatial adjustments. All models were compared to the unadjusted yield. Figure 3 shows the variation in the *CZ* value across different selection intensities and the methods of spatial adjustment. The x-axis is the adjustment method, and the y-axis is the *CZ* coefficient. The median *CZ* across all three selection intensities was 0.82, 0.74, 0.88, and 0.70 for the Moving Mean, Spline, XGB Global and XGB Local, respectively. To put this in perspective, 18% of the lines that were selected by the Moving Mean method were dissimilar from no-adjustment while 30% of the lines that were selected by the XGB Local model were dissimilar from no-adjustment. These results show that utilizing spatial adjustments results in differences in the lines that are selected for advancement within a breeding program. The lowest median *CZ* value observed across all selection intensities was the XGB Local model, meaning that the least amount of overlap between selected lines with no spatial adjustment would be achieved by using this method. However, it should be clarified that spatial adjustments are only needed when there is spatial autocorrelation in yield present within the field. In the absence of yield spatial autocorrelation, the use of unadjusted plot mean in statistical analysis will give the same outcome as spatial adjusted means.

**Figure 3:**
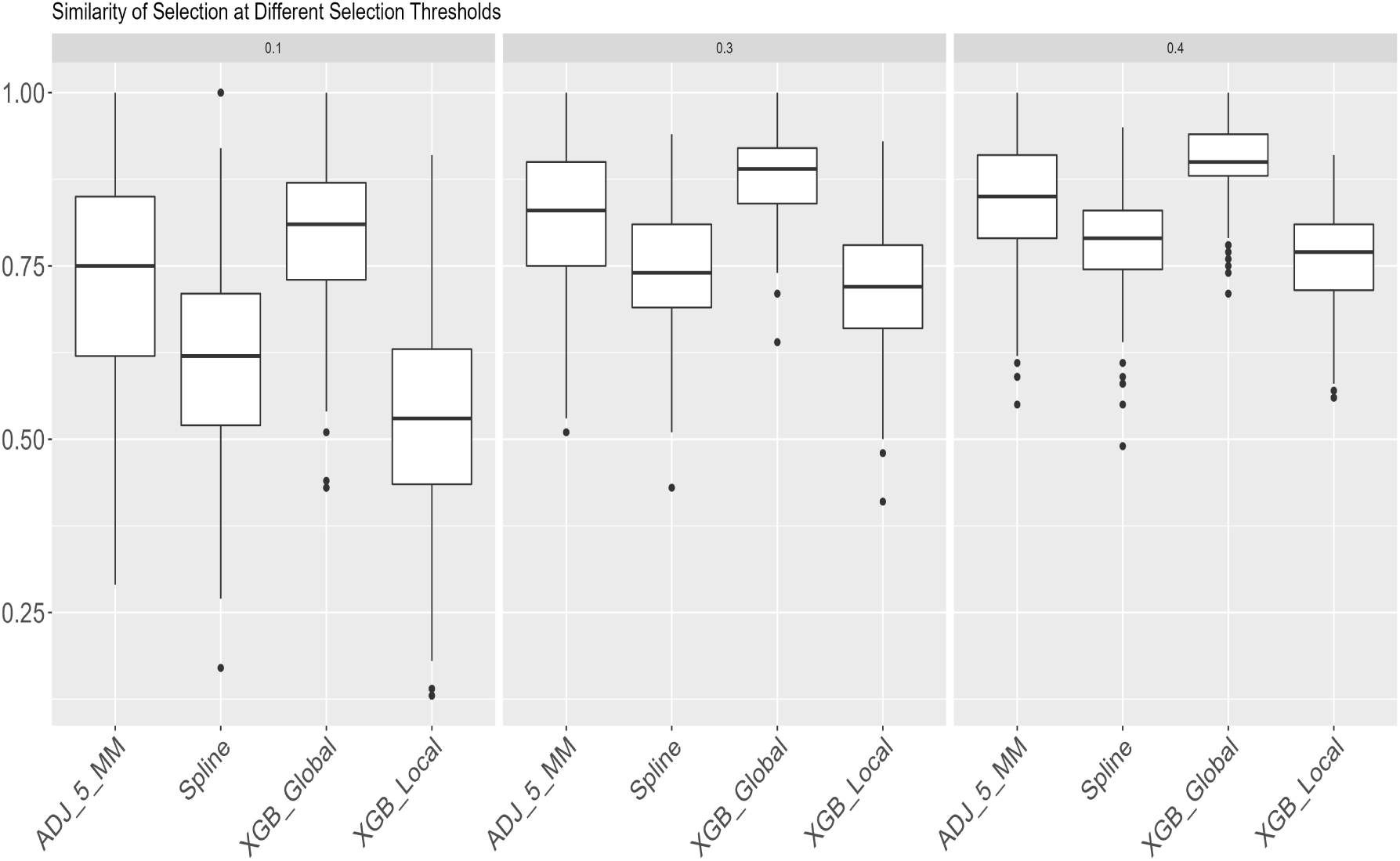
Czekanowski box plots for the different methods. This graph shows the similarity in which lines would be selected between the different models using selection intensities of 0.1, 0.3, and 0.4, in comparison to no spatial adjustment. The XGB Global model exhibited the highest CZ coefficient while XGB Local the lowest. The higher the CZ is, the greater the selection overlapping between the models and the selections made based on no spatial adjustment.

Within breeding trials where lines have similar genetic backgrounds and maturities, it can be expected that there will be similar performance among lines, and because the lines within a trial are randomized it is highly unlikely that there should be evidence of spatial autocorrelation if the environmental effect has been accounted for. The reduction in the spatial trends observed in trials is a good indication that spatial adjustments were effective at removing non-random field effects. Figure 4 shows an example of an experiment before and after the yield has been adjusted using the XGB Local method. The Moran’s I is reduced from 0.58 to 0.09

**Figure 4:**
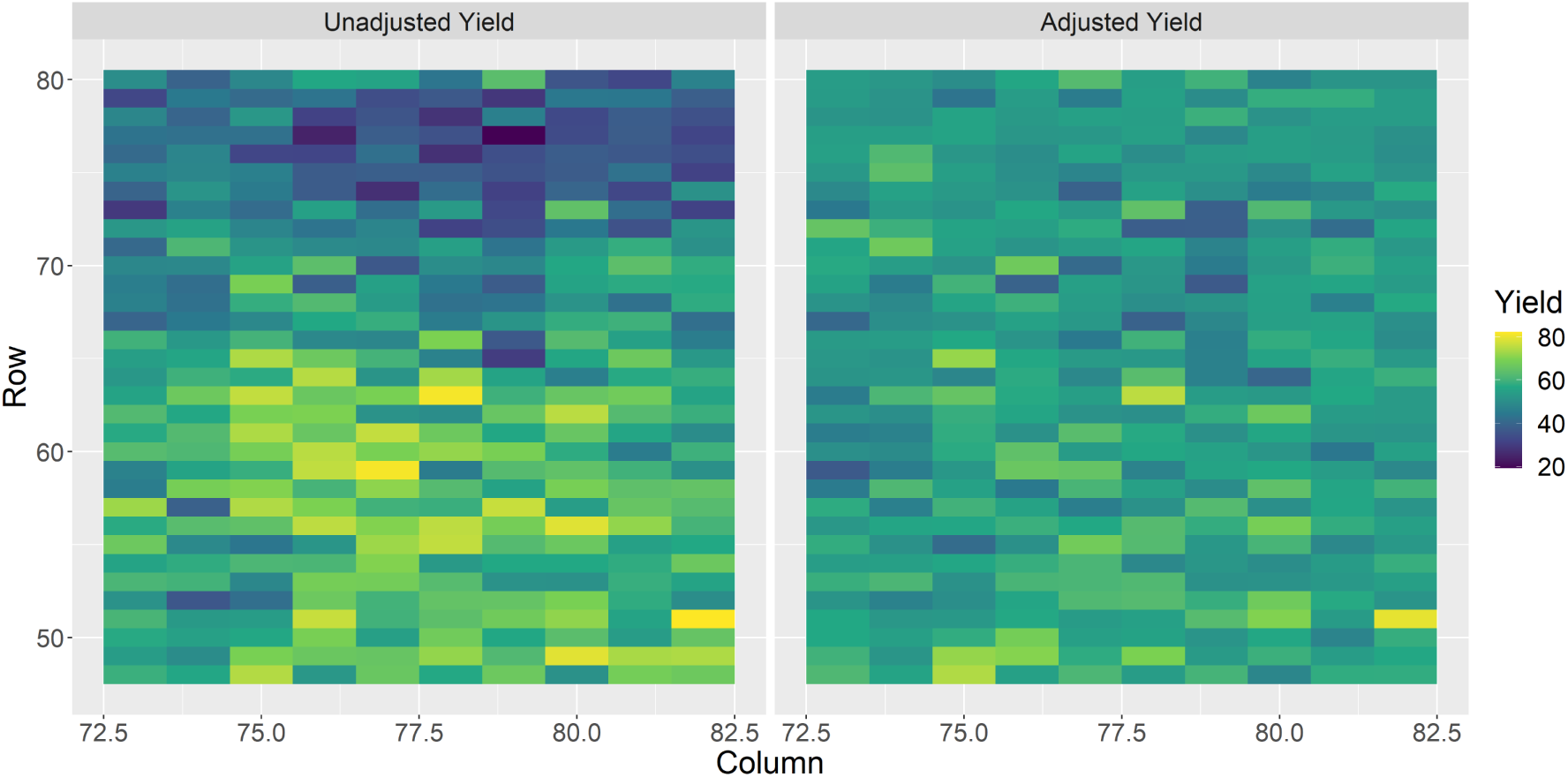
This figure shows the observed yield for both the Unadjusted and Adjusted (XGB Local) yield in an experiment. The Moran’s I for the left image is 0.58, and 0.09 for the right.

Figure 5 shows the range of Moran’s I values across all 238 experiments that were tested. The first two methods, ADJ 5 MM and Spline, are used by breeders for selection decision making. The next two methods are variations of the XGBoost method that used soil data instead of neighbor-based adjustments. The last column labeled as Yield is the Moran’s I values calculated based on no spatial adjustment, and serves as the baseline for all of the comparisons that are made. All methods reduce the spatial autocorrelation within trials, with median Moran’s I values of –0.04, –0.03, 0.04, 0.01, 0.13 for Moving Mean, Spline, XGB Global, XGB Local, and Yield spatial adjustment methods respectively. Figure 5 shows the differing levels of spatial correlation across fields, and provides strong evidence that both traditional neighbor-based adjustments, and soil-based adjustments reduced the overall Moran’s I statistic in the majority of trials. It should be noted that the XGB Local method had a median Moran’s I statistic value (0.01) that was closest to 0 compared to all other methods.

**Figure 5:**
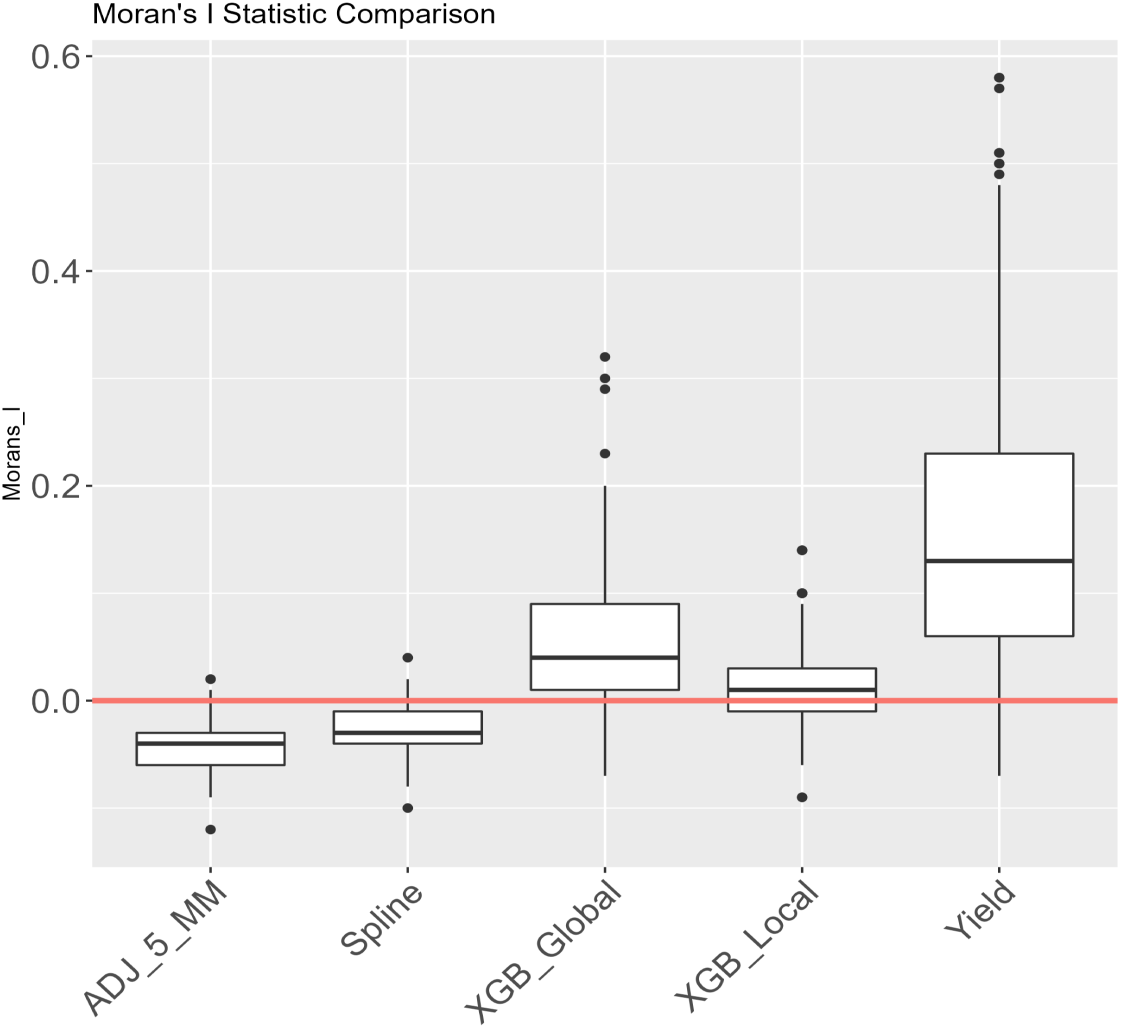
This figure shows the variation of the Moran’s I statistic that was calculated for each experiment. The closer the value is to zero the better evidence there is to suggest that there is a minimal field trend.

## 4 Discussion

Directly modeling the effect of soil on a plot level basis has not been done for plant breeding field trials. The reason for this is that the labor and cost required to collect this type of data is cost prohibitive. As newer technologies arise from the combination of remote sensing and ML this type of data can be obtained at a much lower cost than previously possible [38, 52–54]. Traditional spatial adjustments in field trials have focused on reducing the effect of environmental trends that are evident in most large scale breeding trials. Removing or accounting for the environmental effect of a plot will lead to more accurate selection, and an increase in genetic gain for each breeding program [55, 56]. Soil-based adjustments give breeders additional tools when optimizing for selection of multiple traits [57], such as yield, and biotic stress induced nutrient deficiencies.

Soil fertility recommendations for soybean production in the Midwest have primarily been focused on P, K and pH. Optimal recommended P level is to target *≥* 16 ppm, optimal K levels are recommended to be *≥* 161 ppm [58], and optimal pH range is between 6.5 and 7.5 [8]. Based on these recommendations, 3 of the 9 fields had an average P level that was below optimum, 7 fields had a mean soil pH outside of the optimal zone, and 4 fields had a mean K level below the optimum. It should be noted that these recommendations are not based on maximizing yield, but based on maximizing profitability. Other factors such as CEC have been reported to have an effect on soil nutrient availability and it’s interactions with plant growth and development [59]. These soil nutrients have well established univariate effects on soybean growth and development [9, 10], but when they are combined within multiple interacting complex systems it becomes difficult to quantify the effect of any individual factor. Previous studies have looked at the effects of multiple soil factors and their effects on predicting yield using Random Forest in production fields and found that some of the top-ranking predictor variables were P, K, OM, and pH [60]. Our findings show that these factors had an effect on the quality of the growing conditions of each plot. While feature importance measurements are useful for determining how often a variable is used to split a decision tree, it does not provide any additional insight into the effect that a variable has on a modeled outcome.

The three metrics we used to evaluate the different selection methods examine the utility of different spatial adjustment methods. Neighbor-based adjustments make an assumption that neighboring genotypes should be phenologically similar to reduce the effects of interplot competition [40]. Neighbor-based adjustments are based on linear models, which follow the assumptions that the samples come from the same distribution. If this assumption is no longer valid it can result in poor performance of spatial adjustment methods. This is why experiments usually consist of similar maturity groups and genetic backgrounds, which helps to reduce the interplot competition as genotypes invariably have similar phenological/architectural traits. The XGB Local model had the best metrics for reducing the spatial autocorrelation for seed yield within tests, greatest increase in the mean relative efficiency, and also had the lowest median *CZ* coefficient across all tests and selection thresholds. For caution, it is important to clarify that dissimilar selection decisions stemming from unadjusted or any of the spatial adjustment method require further replicated field trials in the following year to confirm the accuracy of selection of different methods. Since we used an active breeding pipeline for this research, which used Moving Means in A-test in the previous year, we do not have a completely independent set to compare the efficiency of selection across methods. The advantage of using soil features with XGBoost is that the model is based on the feature space of each field where as neighbor-based adjustments utilize only the location space of each dataset. Utilizing the feature space becomes more important when breeders want to grow dissimilar genotypes in the same test. This happens because the nearest neighbor methods have a proclivity for better estimates for similar phenotypes. The other advantage of XGBoost is interpretability of the adjustments giving increased confidence in breeders’ decision making.

The overall results suggest that using a locally trained model using the proposed methodology can increase the efficiency of selection within the breeding program. Interestingly, the local model consistently performed better than global model for applications of spatial adjustments within a breeding program. We hypothesize that this is due to the variation in environments, and that the local models can pick up on environment specific factors. While the global model was not generalized across fields, the non-generalizable models were effective because they were able to determine the local field effects and interactions, whereas the global model became a statistical average of all models and removed the sensitivity that was obtained in the local model that was refined for each individual field.

There are several avenues for integration of ML based spatial adjustment models in other on-going areas of research in crop science. For example, crop modeling has been shown to be an effective tool for parsing out the effects of genotype, environment, and management for crop production [61]. The fusion of better environmental data and increased modeling capabilities, along with HTP methods for genotype specific calibrations, can be used by breeders to make selections for both current and future breeding environments [1, 62]. High resolution soil maps and spatial adjustments can be invaluable for these modeling projects, and it has been shown that spatial adjustments can be useful for increasing genomic prediction model accuracies [63, 64]. Comprehensive weather and genetics data have been used to predict complex phenotypic traits [34, 65, 66], and integration of soil maps and interpretable spatial adjustments can improve the explainability of genotype response. These advantages extend to study of component traits in seed yield and other physiological end-trait predictions [22, 67–69], including for ML based predictions and prescriptive breeding approaches [33, 70].

As previously explained, soil mapping and interpretable spatial adjustments can be useful to study abiotic and biotic stress responses particularly in conjunction with HTP and ML [71–73]. Furthermore, advances in root trait studies [23, 24, 74–76] can be complemented with the use of soil maps and spatial adjustments for a more holistic interpretation of plant response accounting for both above– and below ground traits.

There are four main avenues to further improve this work. (a) The setup of this experiment does not allow us to make causal inferences about the true effect of various soil parameters, but it does open the door for future work where researchers can start to investigate nutrient use efficiency for different genotypes [77], and also the complex interactions of soil, genotypes, weather, and management for optimal plant growth and development. (b) Future work can investigate the use of soil, weather, genotype, and remote sensing data to develop a generalized model for yield prediction. While we included 282 unique populations, it still does not fully represent the entire genetic diversity of the USDA germplasm collection as our testbed program aims to develop elite soybean varieties. We controlled for variations in genetic background by centering populations around their mean values, but more work looking into more crops should be done to determine the effectiveness of this methodology. More recent work that studied the effect of including genetic relationships in spatial adjustment methods has shown that including SNP and pedigree information can help to improve spatial adjustments; however, this was based on simulated data [78]. As breeders continue to have more tools at their disposal modeling the genetic and environment factors could help to make more informed selections. This can remove non-genetic effects when evaluating lines, and also start to investigate the genetic components that effect soil nutrient use efficiency and help to create more prescriptive cultivars across regions [70]. (c) The analysis methods presented in our study were not used to compare performance in the following years, to see how different selection methods performed. Future work should investigate the repeatability of methods across years and locations. (d) Variable soil fertility was observed across all testing sites, but the locations that were tested are generally considered to be high yielding environments. Future work investigating the utility of soil mapping in lower productive soil is needed to see the utility of this method in a wider range of environments.

## 5 Conclusion and Future work

We demonstrate the usefulness of spatial adjustment models to improve the selection efficiency in plant breeding programs by removing environmental trends that can bias selection decision making. While spatial adjustments do not give any advantage over unadjusted plot seed yield based analysis when no field variation exist, such a scenario is extremely rare. Therefore, unreplicated trials will benefit from the use of spatial adjustment models. In this paper, we demonstrate the usefulness of a ML method for spatial adjustment in plot experiments. This method models soil fertility trends within a field, and allows breeders to increase their knowledge about the quality of the growing conditions of each plot within a field trial. Compared to unadjusted, Moving Mean and Spline methods, the XGB Local model had the greatest increase in the mean relative efficiency, the lowest median *CZ* coefficient across all tests and selection thresholds, and showed a consistent ability to reduce field gradient effects on the yield of plots. The XGBoost method we demonstrate is based on soil features, which becomes more important when breeders want to grow genetically diverse and phenologically dissimilar genotypes in the same test where the soil map indicates variability in nutrients, soil texture properties, pH, CEC, and OM. XGBoost provides interpretability of the adjustments that can increase breeders’ confidence in the application of spatial adjustment methods. Additionally, soil-based adjustments provide opportunities to select for genotypes that respond to soil features, for example, nutrient deficiency response.

## Acknowledgments

General

We thank staff and student members of Singh Soybean group at ISU, particularly Brian Scott, Will Doepke, Jennifer Hicks, and Ryan Dunn for their assistance with field experiments and phenotyping. We thank Sarah Jones and Ashlyn Rairdin for help in editing the manuscript. We thank Yones Khaledian and Caner Ferhatoglu for generating the soil maps, and members of the Geospatial Laboratory for Soil Informatics group for their assistance in soil core sampling.

## Author contributions

A.K.S and M.E.C conceived the research; M.E.C. and A.K.S. performed experiments and data collection; B.A.M. contributed to the interpretation of soil data; P.M.D. contributed to the interpretation of statistical results; M.E.C. and L.G.R. built machine learning models and interpreted the results with inputs from S.S., B.G., and A.K.S.; M.E.C. and L.G.R. prepared the first draft with A.K.S. and S.S. All authors contributed in the development of the manuscript, and reviewed the manuscript. M.E.C. and L.G.R., as co-first authors, made equal contributions to the paper.

## Data Availability

Data and Codes will be made available publicly after acceptance of the paper through corresponding authors’ GitHub pages.

## Supplementary Material

**Figure 6:**
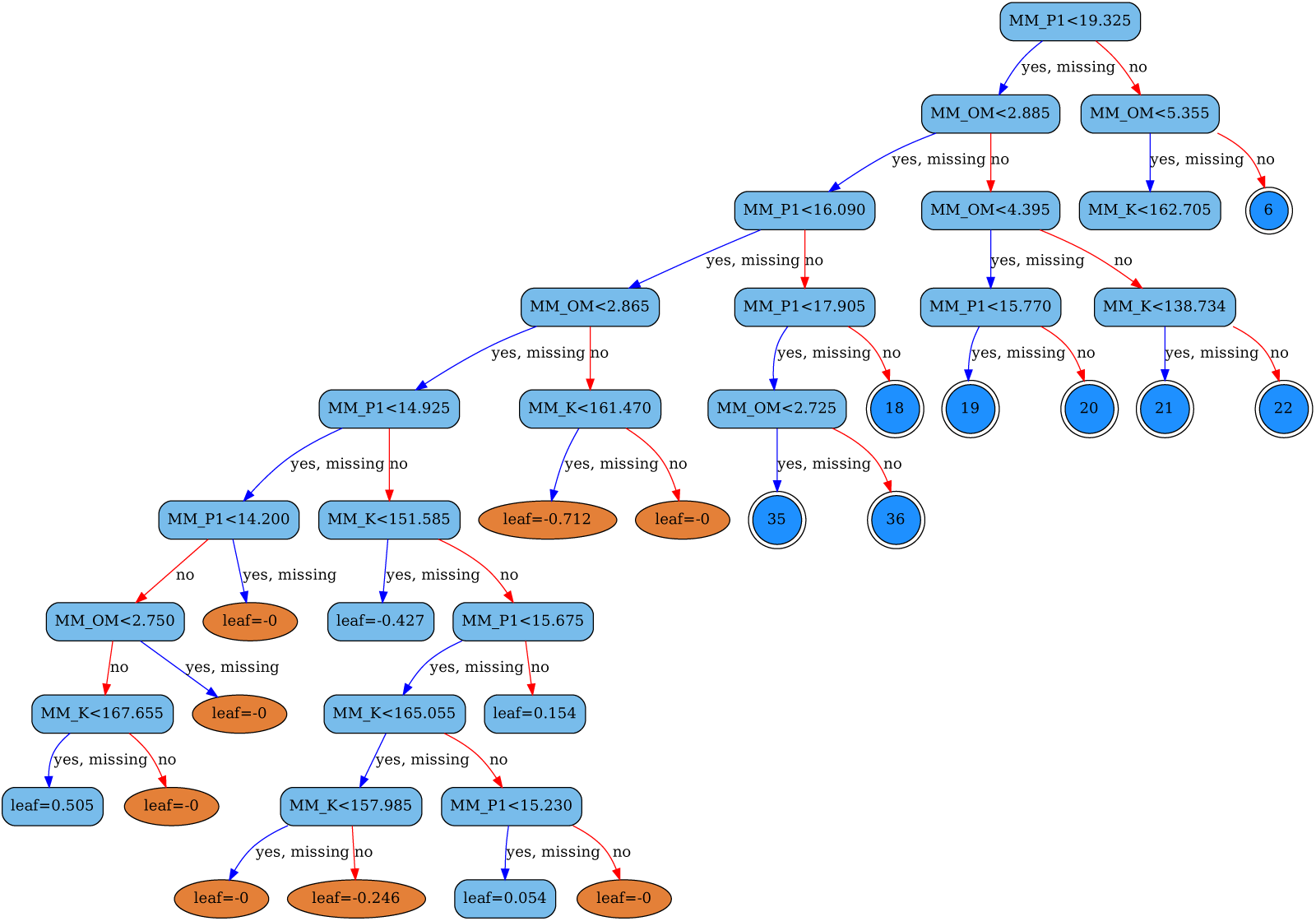
An example of a trimmed XGBoost tree model stopping at a depth of eight. The circles represent other branches no shown here, the ellipses are the leaf, and boxes are the splitting decision nodes.

**Figure 7:**
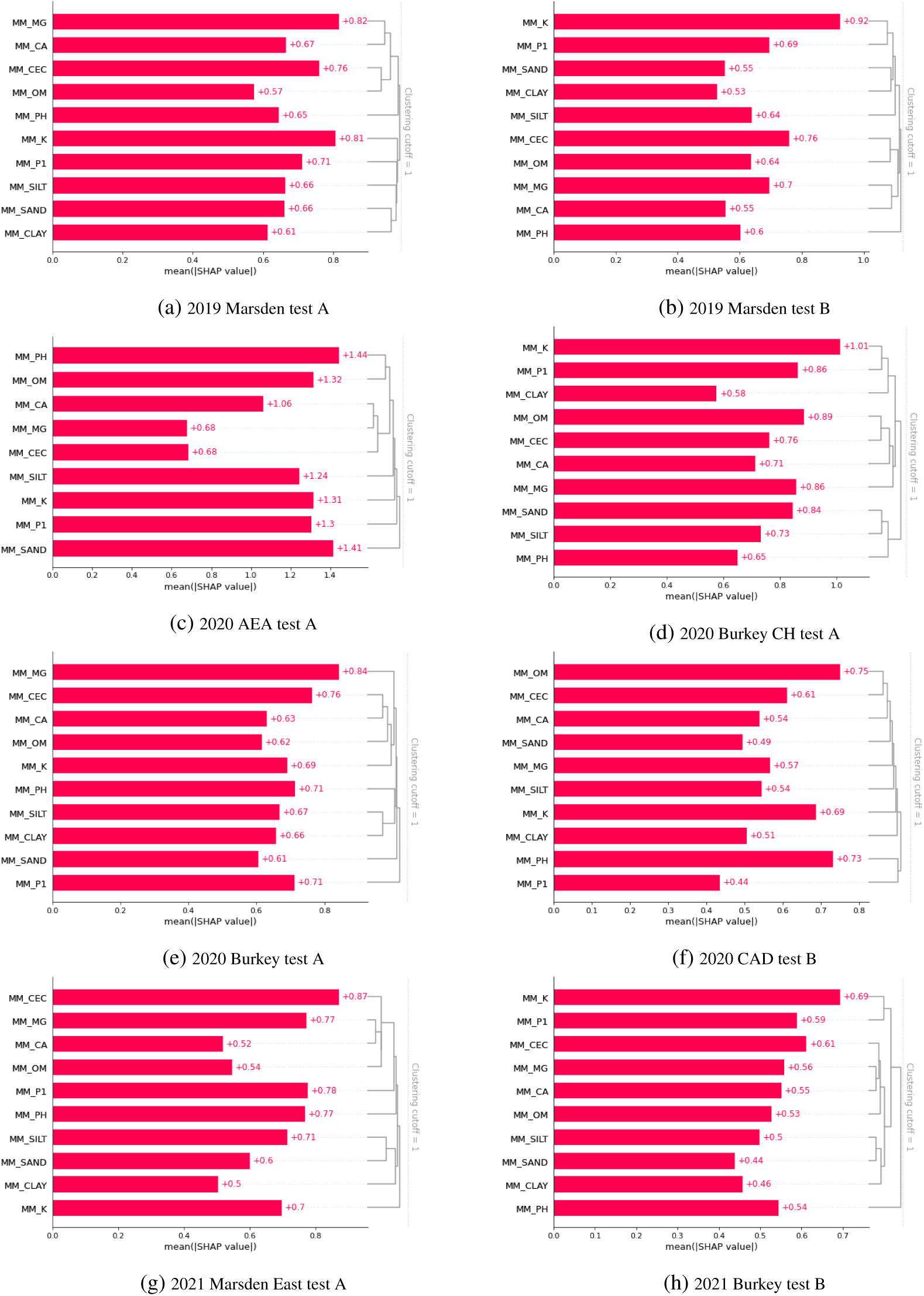
XGBoost feature importance plots. The mean(—SHAP Value—) is a weighted measurement based on the number of times a feature was used by the model to split the data across all trees. The splitting is done as function of information gain. The higher the score the more important the the feature is for the given model. The dendrogram on the right of the plots represent how the model clusters the different features using average euclidean distance and threshold of one.

**Figure 8:**
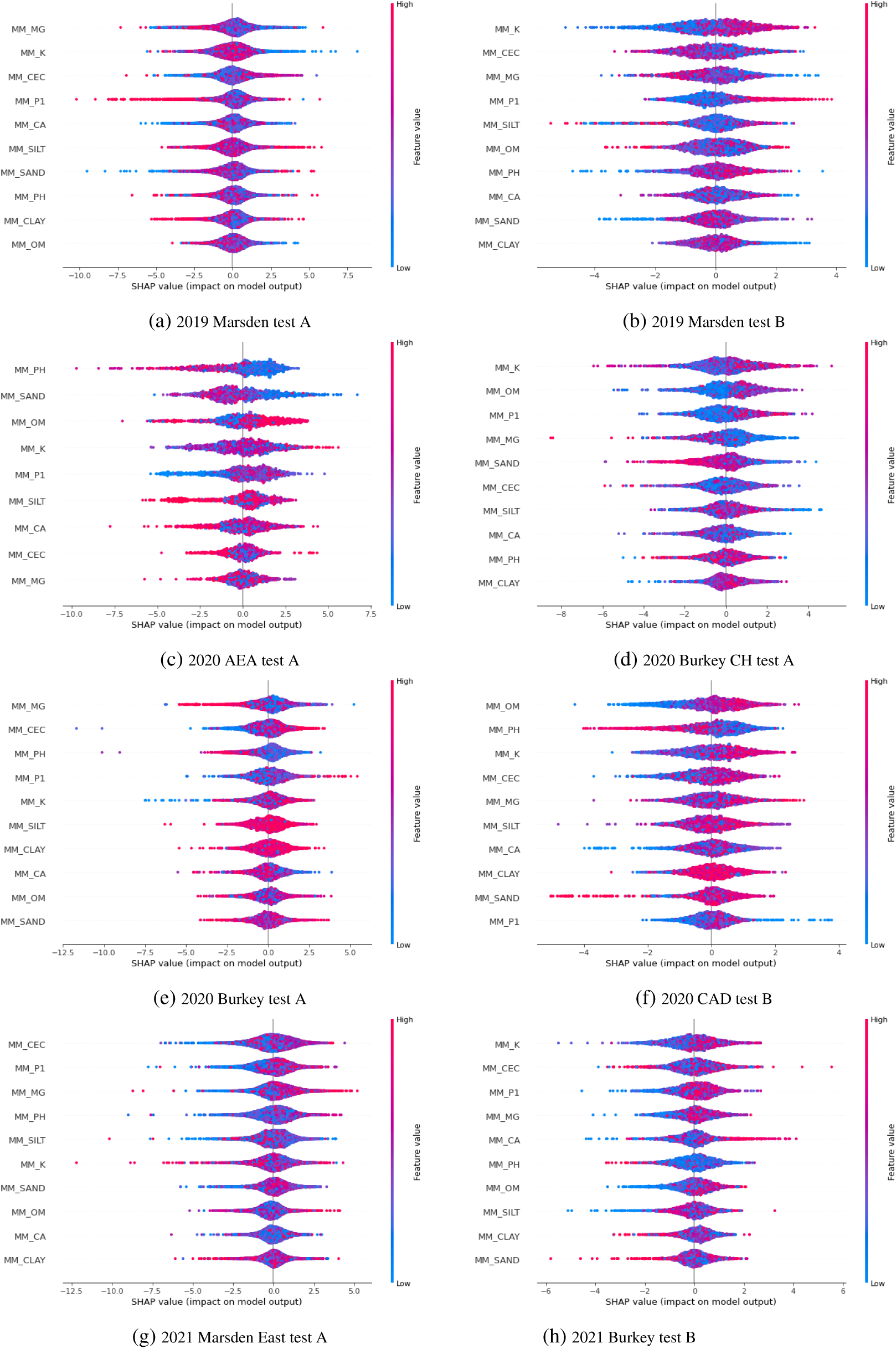
Explanatory Shapley Values summary plots.

**Figure 9:**
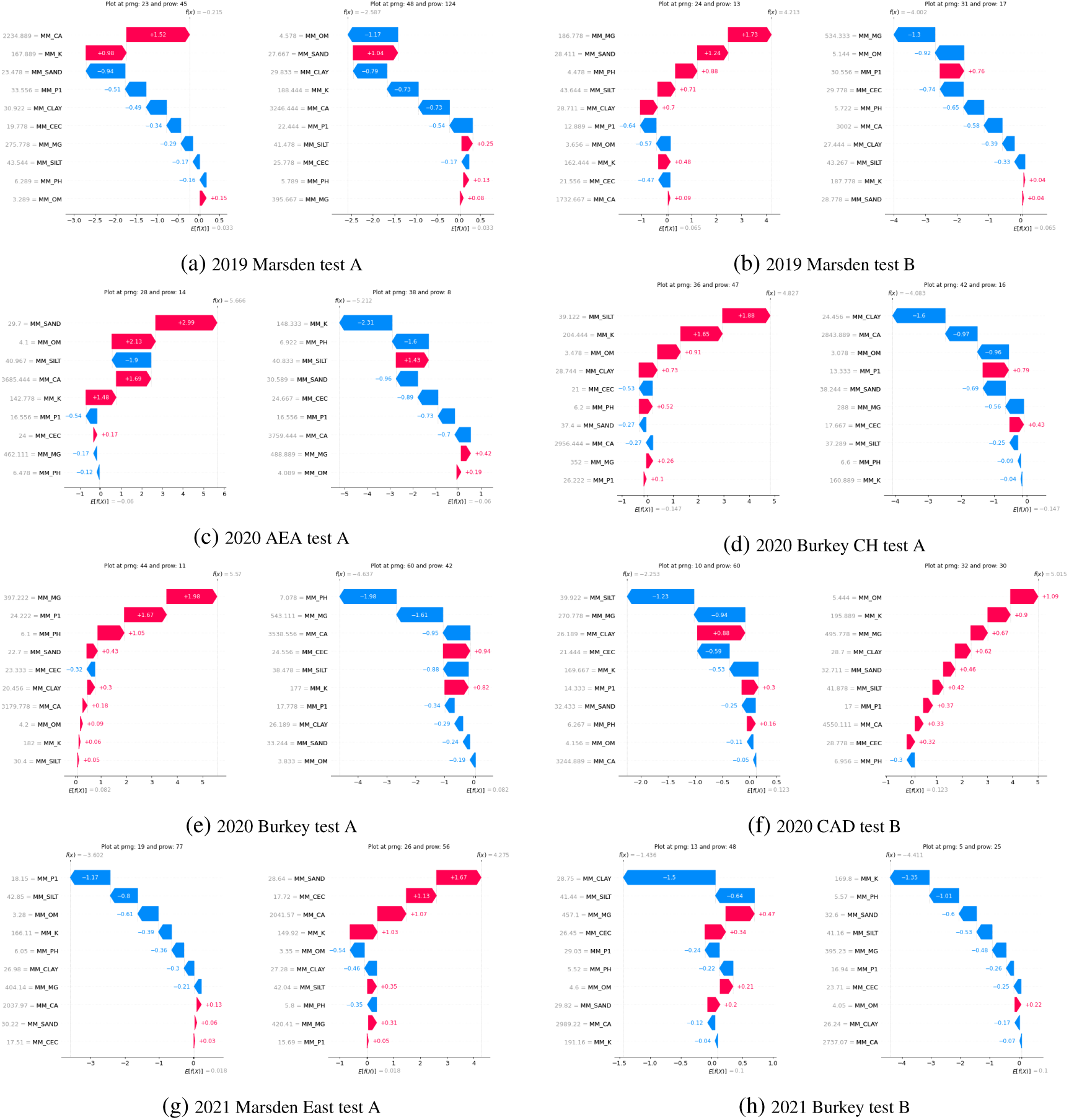
Explanatory Waterfall charts for genotype IA2102 across the nine experiments. There are two Waterfall per subplot, each showing how the feature values contributed to the final predicted yield at the given range and row field coordinates. Feature measurements are to the left of the feature names, while the Shapley values are within the arrow bar. The predicted means for the model are on the x-axis, and the predictions for the plot, shown on top, are obtained by adding and subtracting the respective Shapley values.

